## Supplementary figures and images for "A non-conducting role of the Ca_v_1.4 Ca^2+^ channel drives homeostatic plasticity at the cone photoreceptor synapse"

### Figure 2-Figure Supplement 1

**Figure 2- Figure Supplement 1**

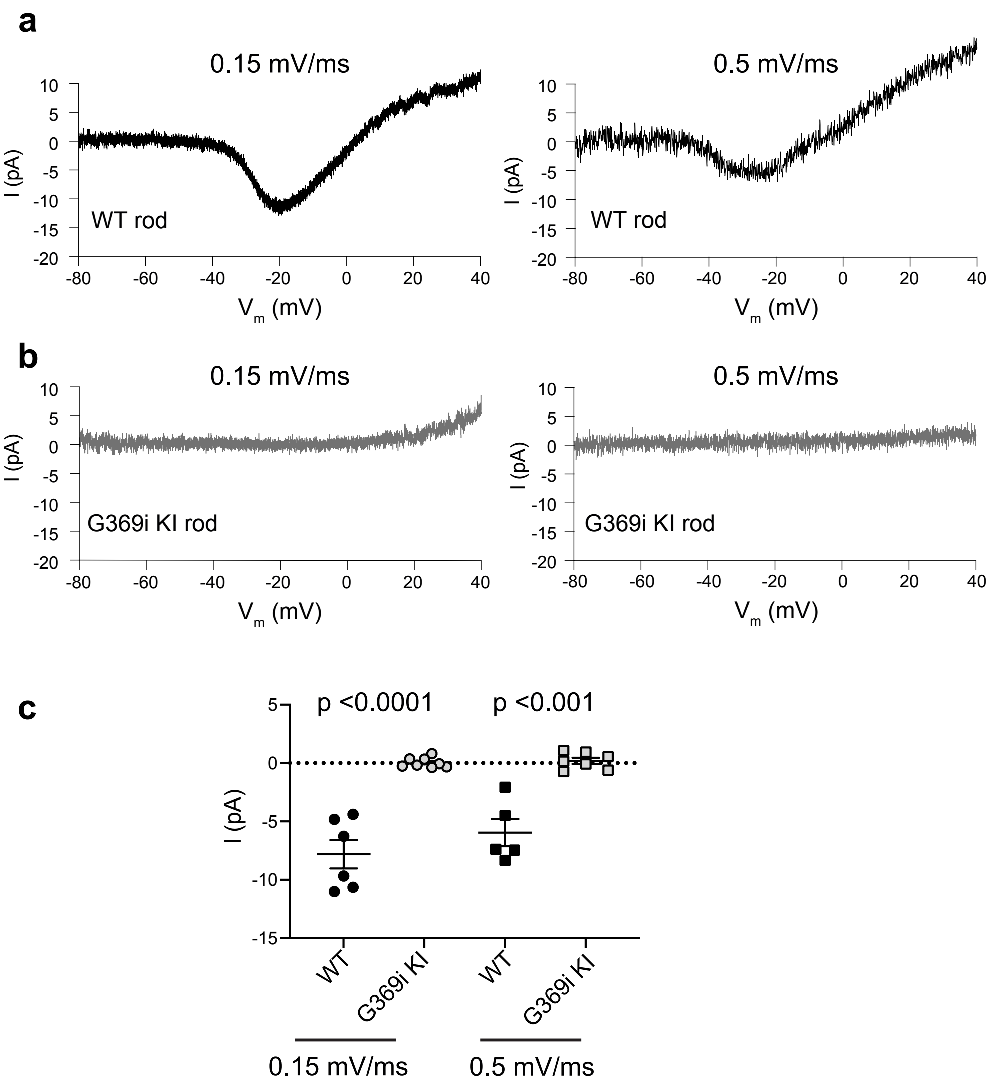

### Figure 2-Figure Supplement 3

Figure 2- Figure Supplement 3.

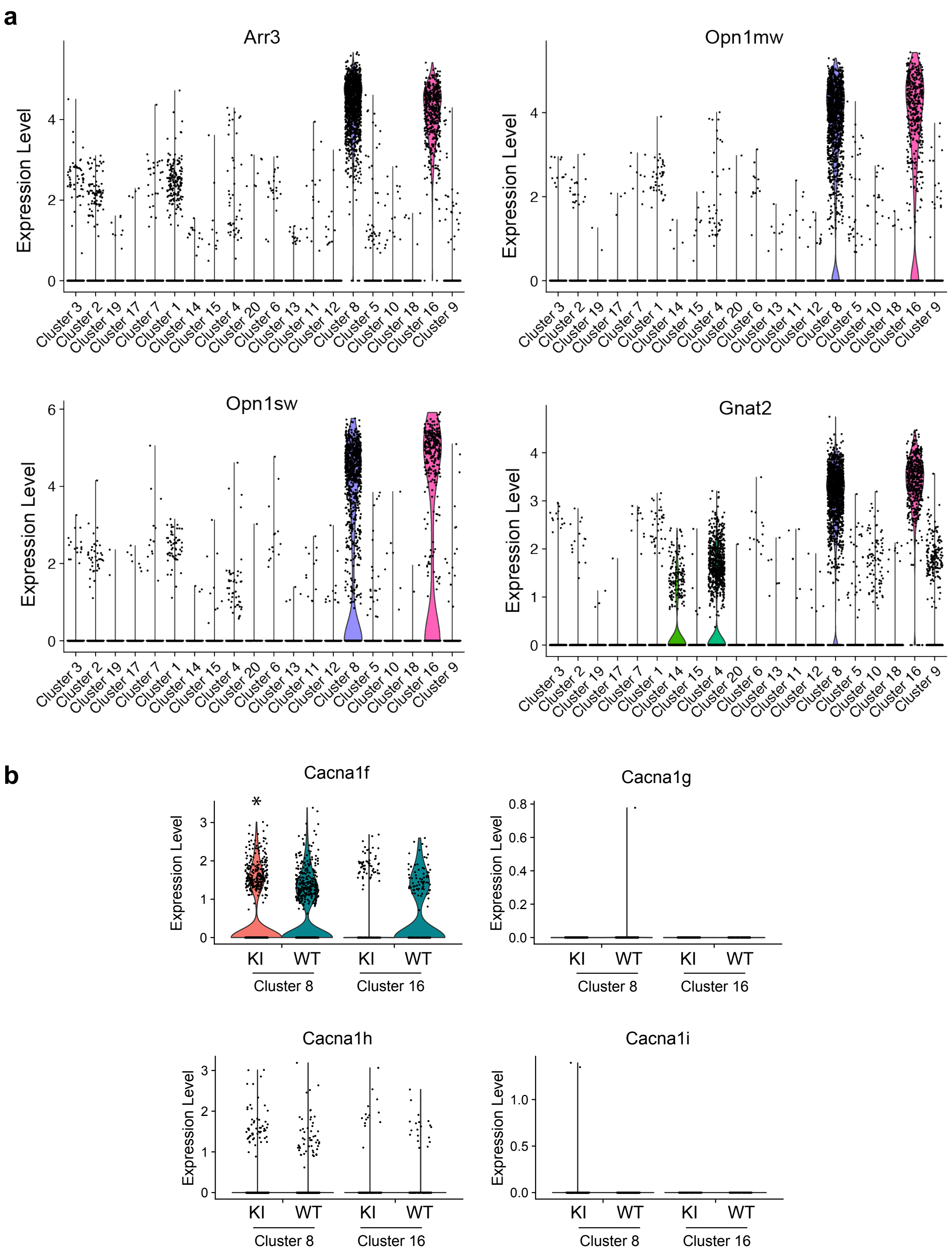

### Figure 3-Figure Supplement 2

Figure 3- Figure Supplement 2

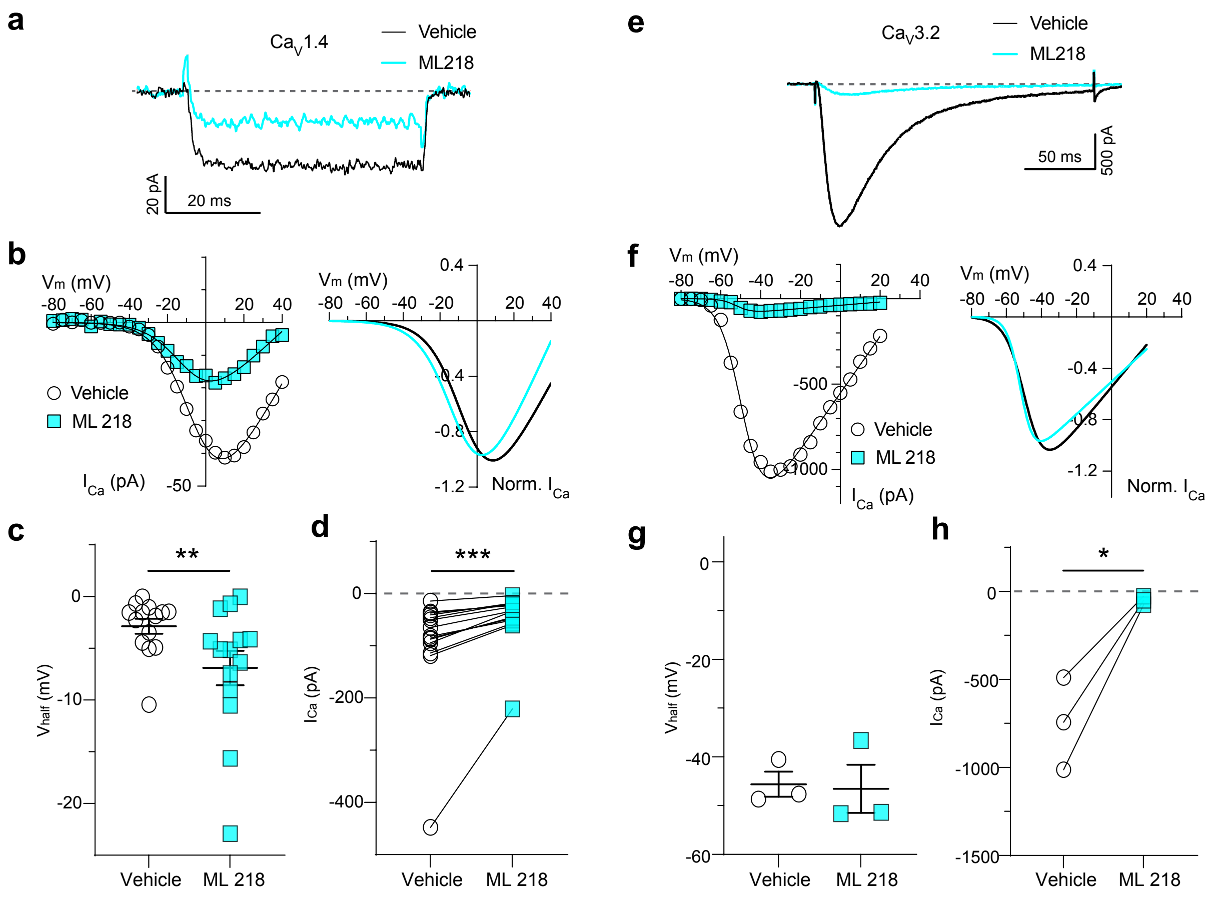
